# Population-specific responses to developmental temperature in the arboviral vector *Aedes albopictus*: implications for climate change

**DOI:** 10.1101/2023.10.06.561151

**Authors:** Martina Carlassara, Ayda Khorramnejad, Helen Oker, Romina Bahrami, Alejandro Nabor Lozada-Chávez, Maria Vittoria Mancini, Mélanie J.A. Body, Chloé Lahondère, Mariangela Bonizzoni

## Abstract

The increase of environmental temperature due to current global warming is not only favoring the expansion of the distribution range of many insect species, but it is also changing their phenology. Insect phenology is tightly linked to developmental timing, which is regulated by environmental temperatures. However, the degree to which effects of developmental temperatures extend across developmental stages and their inter-stage relationships have not been thoroughly quantified in mosquitoes. Here, we used the mosquito *Aedes albopictus*, which is an aggressive invasive species and an arboviral vector to study how developmental temperature influences fitness across developmental stages, thermal traits, energy reserves, transcriptome, and *Wolbachia* prevalence in populations from either temperate or tropical regions. We show that hatchability, larval and pupal viability, and developmental speed are strongly influenced by temperature and these effects extend to wing length, body mass, longevity, content of water, protein and lipids in adults, in a population-specific manner. On the contrary, neither adult thermal preference nor heat resistance significantly change with temperature. Development at 18°C revealed to be a limiting factor for the proliferation of *Wolbachia* in adults and transcriptome analysis showed enrichment for functions linked to stress responses (i.e. cuticle proteins and chitin, cytochrome p450, and heat shock proteins) in mosquitoes reared at both 18°C and 32°C. Our data showed an overall reduced vector fitness performance when mosquitoes were reared at 32°C, and the absence of isomorphy in the relationship between developmental stages and temperature in the temperate population. Altogether these results have important implications for reliable model projections of the invasion potentials of *Ae. albopictus* and its epidemiological impact.

## INTRODUCTION

Thermal conditions experienced during development have immediate and long-lasting effects in animals, in particular in poikilothermic organisms with an holometabolous life cycle like mosquitoes (Mirth et al., 2021; Müller et al., 2016; Nussey et al., 2007; Van De Pol et al., 2006; Vasudeva et al., 2014). For example, at lower temperature, mosquitoes are expected to grow larger because of slower growth rates of immature stages (Atkinson, 1994). An indirect outcome of this phenomenon is that traits influenced by size, such as fecundity and adult longevity (Nørgaard et al., 2022) may be in turn affected. Developmental time, fecundity, and adult longevity are directly related to the transmission potential of a mosquito vector. A quicker development and a higher fecundity may increase the adult density and reproductive success. Moreover, older vectors are more likely to have been exposed to an infectious blood meal and have survived the extrinsic incubation period of any pathogen they carry (Cator et al., 2020). Consequently, developmental temperature plays a major role in the fitness performance of a vector mosquito and should be investigated in light of current global warming that is favouring advances in seasonal phenology (Parmesan, 2006). However, the extent to which thermal conditions experienced during development affects not only juvenile, but also adult stages, have not been thoroughly quantified in mosquitoes. Additionally, in the context of invasive mosquito species, which may experience high levels of thermal variation, understanding the relationship between developmental temperature and fitness across developmental stages is critical. Whether this relationship differs among geographic populations could possibly lead to the identification of not only the “most adaptable” and invasive population(s), but also of the most vulnerable life-stage to tackle with control strategies considering current global climate changes.

The Asian tiger mosquito *Aedes albopictus* is a highly invasive mosquito of prime public-health relevance due to its vector competence for arboviruses such as dengue, Zika and chikungunya (Bonizzoni et al., 2013; Manni et al., 2017; Pichler et al., 2019; Vavassori et al., 2022). In the past 50 years, this species has conquered every continent, except Antarctica, from its native home range in Asia and from islands of the Indian Ocean and the Pacific, which it had colonised during the spice trade (17th-18th century) (Delatte et al., 2011; Ducheyne et al., 2018). Remarkably, temperate populations of *Ae. albopictus* undergo photoperiodic diapause and overwinter predominantly as eggs, suggesting profound physiological adaptations to cooler climates, whereas tropical populations do not. The physiological and molecular mechanisms of photoperiodic diapause have been well studied in *Ae. albopictus* (Armbruster, 2016; Boyle et al., 2021). However, we lack data on traits that have been shown important for arboviral transmission such as larval viability, developmental time, adult thermal tolerance and adult longevity, all of which can be affected by thermal conditions experienced during development (Folguera et al., 2010; Mordecai et al., 2019)

Amongst the most recent and eco-friendly approaches to control *Ae. albopictus* populations is the application of the symbiont *Wolbachia* to drive cytoplasmic incompatibility (CI) or control arboviral infection (Ant et al., 2018; Moreira et al., 2009; Walker et al., 2011). CI is a phenomenon whereby the mating of a male carrying a particular *Wolbachia* strain with a female lacking that strain, or carrying an incompatible *Wolbachia* strain, results in no progeny (Hoffmann & Turelli, 1997). *Aedes albopictus* naturally harbours two strains of *Wolbachia*, *w*AlbA and *w*AlbB, which can both induce CI (Kittayapong et al., 2002; Sinkins et al., 1995). Additionally, when females infected with either *w*AlbA or *w*AlbB are crossed with superinfected males, unidirectional CI is obtained (Yang et al., 2022). While in nature *Ae. albopictus* mosquitoes tend to show superinfections, the infection frequencies and densities of the native strains might vary significantly across populations (Yang et al., 2022). Among other factors, environmental temperature (T*_a_*) has been recognised as an important abiotic modulator of density, CI penetrance, and maternal transmission fidelity, having variable degree of intensity among different *Wolbachia* strains (Mancini et al., 2020; Ross et al., 2017). Moreover, the impact of developmental temperatures on *Wolbachia*-conferred antiviral protection in natural and transinfected hosts was also demonstrated (Chrostek et al., 2021; Mancini et al., 2021).

In this framework, we investigated how different developmental temperatures affect the fitness of *Ae. albopictus* mosquito larvae of a tropical and a temperate population and how these changes extend to the adult stage, including their fitness and physiology and the density of *Wolbachia*. We further compared the transcriptome of both larvae and adults of the two populations reared under different thermal conditions to link fitness and physiological processes at the molecular level. Chosen developmental temperatures include 18, 28 and 32°C. These temperatures are ecologically relevant: 18°C represents the average daytime temperature registered in northern Italy between April-May 2021 (source: World-weather, https://world-weather.info/forecast/italy/crema/may-2021/), when mosquitoes started to emerge from the winter season; 32°C is the average peak of heat that occurred in the same region during August 2021 (source: World-weather, https://world-weather.info/forecast/italy/crema/august-2021/), while 28°C is the standard *Ae. albopictus* rearing condition (Palatini et al., 2020).

We show extensive population-specific effects on the fitness and energy reserve of mosquitoes reared under different thermal regimes, whereas we did not observe significant thermal-related changes in adult thermal preference or heat resistance. In both populations, the fitness performance was reduced when mosquitoes were reared at 32°C. Development at 18°C revealed to be a limiting factor for the proliferation of *Wolbachia* in adults and transcriptome analysis showed enrichment for functions linked to stress responses (i.e. cuticle proteins and chitin, cytochrome p450 and heat shock proteins) in mosquitoes reared at both 18°C and 32°C. We further demonstrate an absence of isomorphy in the relationship between developmental stages and temperature in the temperate, but not the tropical population (Folguera et al., 2010; Jarošík et al., 2004) which has important and practical applications as developmental proportionality limits the evolution of life-history traits in response to thermal changes. We discuss our results in relation to the invasion history of *Ae. albopictus* and its future potentials.

## MATERIAL AND METHODS

### Mosquito populations

In this study we used two *Ae. albopictus* population: the tropical Foshan (Fo) and temperate Crema (Cr) population. Foshan was established in the early 1980s from Foshan (China) and is the genome-reference *Ae. albopictus* strain (Chen et al., 2015; Palatini et al., 2020). Foshan has been maintained at the University of Pavia since 2013, as previously described (Palatini et al., 2017). Crema was derived from larvae collected in the city of Crema (Italy) in September 2017. Since establishment, the two populations have been maintained in parallel at 28°C and 70-80% relative humidity, with a light/dark cycle of 12 h. Larvae are reared in plastic containers (BugDorm 19×19×6cm) at a controlled density (about 200 larvae in 1 L of water) to avoid competition for food, which is provided daily in the form of fish food (Tetra Goldfish Gold Colour). Adults are fed with cotton soaked in 20% sugar as a carbohydrate source. Adult females are offered commercial defibrinated mutton blood (Biolife Italiana) using a Hemotek feeding apparatus.

### Thermal regimes, assessment of fitness and thermal traits

For both Fo and Cr, a total of twenty-four groups, consisting each of 100 eggs, were placed in plastic containers (17 x 6.5 x 12 cm), with 200 ml autoclaved water, and hatched under different thermal regimes. A set of eight groups of eggs was hatched and reared until adult emergence at 18°C; a set of eight groups of eggs was hatched and reared until adult emergence at 32°C; an additional set was kept at standard 28°C. To avoid confounding effects of humidity and photoperiod cycles, these parameters were maintained equal across thermal regimes. While in the wild, mosquitoes are exposed to fluctuating temperatures, having a constant developmental temperature allows to isolate the effect of a unique experimental variable and generate baseline data for future, more complex analyses (Sinclair et al., 2016).

In each tray, emerging larvae were counted daily to assess egg hatching rate and hatching time. Larval viability was calculated as the percentage of larvae becoming pupae. Larval developmental time (LDT) was measured until pupation. Pupal viability and pupal developmental time (PDT) were measured by counting emerging adults and subtracting LDT to the egg-to-adult developmental time. At emergence, adults were sexed, and a sample of 30 females was used to determine wing length as a proxy for adult size (Robert & Olson, 1989). The right wing was dissected and measured from the axial incision to the apical margin, excluding the fringe of the scales (Figure S1). Measurements were carried out under an inverted microscope (Olympus CKX53) using the software cellSens Standard (Olympus). For each thermal regime, we also assessed adult longevity, thermal preference (*Tp*), namely the temperature at which mosquitoes prefer to lay, and heat resistance as knock-down temperature (*KDT*), namely the temperature and time when the mosquito stopped to move (Van Heerwaarden & Sgrò, 2011). Briefly, for each strain, 500 eggs were hatched at 18°C, 28°C or 32°C, and adult survival was monitored individually until death for males and females separately. To measure *Tp,* 5-9 day-old mosquitoes were collected and isolated by cooling down individuals on the day of emergence in a 4°C fridge. Mosquitoes were released at the centre of a custom-built thermal gradient connected to a cool water bath on one side and to a warm water bath on the other, as in Reinhold et al (2022). These were set at a specific temperature to create a continuous gradient along the aluminium plate. The gradient of temperatures between 15.3-41.5°C had increments of 0.79°C ± 0.285 from one side to the other, with the centre having a temperature of 27.9°C±0.85. After a 5 minutes adaptation time, mosquitoes were monitored for 30 minutes until they settled on a resting spot corresponding to a specific temperature, which was defined as the mosquito *Tp*. A maximum of ten sugar-fed individuals of the same sex, population and thermal regime were released on the thermal gradient each time. A total of eight replicates of ten mosquitoes each were conducted for each sex and population of mosquitoes reared at each of the tested thermal regimes. KDT was determined for individual mosquitoes using a custom-made device. Briefly, an aluminium plate with nine wells, each holding a single 5-9 day-old mosquito, was set at 25°C using a Peltier. Temperature was then increased with increments of 0.5°C / min, until 50°C was reached. A camera (Logitech C922 Pro) connected to a computer was placed above the device to monitor mosquito behaviour and determine mosquito KDT. A total of five replicates of nine mosquitoes each for sex, strain and thermal regime were run. Before the KDT experiments, mosquitoes were collected as described for *Tp*. For these experiments, mosquitoes were fed exclusively with 20% sugar, to avoid any bias due to blood feeding and digestion in longevity, *Tp* and KDT measurements (Camargo et al., 2020; Muturi et al., 2021; Nag et al., 2021).

### Colorimetric assay

We used a colorimetry protocol modified after Foray et al., (2012) to assess protein, glycogen, lipids and triglyceride contents. Total sugars were not measured since mosquitoes were fed with sugar. A total of 50 mosquitoes were processed for each sex, population, and temperature condition. Mosquitoes were collected and isolated by cooling down individuals on the day of emergence in a 4°C fridge. Individuals were then weighed using a microbalance and stored in individual tubes at -70°C until processing. An aqueous lysis buffer (pH 7.4) was prepared with the following composition: 100 mM potassium dihydrogen phosphate (Sigma-Aldrich, #7778-77-0), 1 mM dithiothreitol (Sigma-Aldrich, #3483-12-3), 1 mM ethylenediaminetetraacetic acid (Sigma-Aldrich, #60-00-4) in water. One hundred and eighty μL of the aqueous lysis buffer were added to each sample before grinding using polypropylene pellet pestles (KIMBLE, #749521-0500) for approximately 30 seconds. The homogenates were vortexed and then centrifuged at 4°C for 15 minutes at 2,000 rpm.

#### Total protein quantification

For the quantification of total proteins, 5 μL of each bovine serum albumin (Fisher Scientific, #9048-46-8) standard (0, 0.2, 0.4, 0.6, 0.8, 1.0 μg/μL) was transferred to a 96-well borosilicate plate. Then, 2.5 μL of each sample’s supernatant and 2.5 μL of aqueous lysis buffer were added to the individual wells. Two hundred and fifty μL of Bradford micro-assay reagent (Sigma-Aldrich, #B6916) was added to each well, and the plate was incubated in the dark at room temperature for 5 minutes. The plate was then placed in the microplate reader (Accuris SmartReader 96) and shaken at low speed for 5 seconds within the instrument and read at 595 nm. Absorbance values were recorded and used to calculate protein concentration in mosquito samples.

After quantification of protein, the samples were then divided into two tubes: a stock solution and a glycogen pellet. To accomplish this, 2.5 μL of aqueous lysis buffer, 20 μL of sodium sulphate solution (20% sodium sulphate (Sigma-Aldrich, #7757-82-6) in water), and 1500 μL of chloroform-methanol solution (1:2 ratio of chloroform (Sigma-Aldrich, #67-66-3) to methanol (Sigma-Aldrich, #67-56-1) were added into each sample tube. The tubes were vortexed and then centrifuged at 4°C for 15 minutes at 2,000 rpm. Following centrifugation, the supernatant was transferred into a separate 2 mL centrifuge tube and will be referred to as the stock solution. The pellet left in the original tube was then used for glycogen quantification.

#### Glycogen quantification

For the quantification of glycogen, the pellet was initially washed with 400 μL of 80% methanol diluted in water, vortexed, and then centrifuged at 4°C for 5 min at 15,000 rpm. The supernatant was removed, and the washing step was repeated once more. Glucose (Sigma-Aldrich, #50-99-7) standards were made by adding 25 μL of each standard (0, 0.5, 1.0, 1.5, 2.0, 2.5, 3.0 μg/μL) into individual tubes and allowing the solvent to fully evaporate (approximately 36 hours at room temperature). After supernatant evaporation, 1 mL of anthrone solution (0.142% w/v anthrone (Sigma Aldrich, #90-44-8) in 70% sulfuric acid (Sigma-Aldrich, #7664-93-9) diluted in water was added to each sample and standard and then vortexed. The tubes were then incubated at 90°C for 15 minutes and vortexed halfway through the incubation period. The tubes were then placed on ice for at least 5 minutes and then 250 μL of each sample and standard was transferred into a 96-well borosilicate plate. Next the plate was read at 625 nm. Absorbance values were recorded and used to calculate glycogen concentration in mosquito samples.

#### Total lipids quantification

For the quantification of total lipids, the stock solution of each sample was vortexed and then 150 μL of each sample was transferred to a 96-well plate. Five μL of each triolein (Sigma-Aldrich, #122-32-7) standard concentrations (0, 0.5, 1.0, 1.5, 2.0, 2.5, 3.0 μg/μL) was also transferred to the plate. The plate was incubated at 90°C until the solvent fully evaporated. 10 μL of 98% sulfuric acid was then added to each sample and standard and the plate was once again incubated at 90°C for 2 minutes. The plate was then placed on ice for 5 minutes and 190 μL of vanillin solution (0.120% w/v vanillin (Fisher Scientific 121-33-5) in 68% orthophosphoric acid (Sigma-Aldrich 7664-38-2) diluted in water) was added to each well. The plate was then shaken at low speed in the plate reader for 15 minutes and read at 525 nm. Absorbance values were recorded and used to calculate total lipid concentration in mosquito samples.

#### Triglycerides quantification

For the quantification of triglycerides, each sample’s stock solution was vortexed and then 500 μL of stock solution was transferred to a new 2 mL centrifuge tube. Each tube was incubated at 90°C until the solvent completely evaporated. The tubes were removed from the heat and 1 mL of chloroform and 200 mg of anhydrous silicic acid (Sigma-Aldrich 10279-57-9) was added to each tube. The tubes were centrifuged at 4°C for 10 minutes at 2,000 rpm. 400 μL of each sample’s supernatant was transferred into individual wells of a 96-well plate. 5 μL of each triolein standard (0, 0.5, 1.0, 1.5, 2.0, 2.5, 3.0 μg/μL) was also transferred to the plate. The plate was incubated at 90°C until the solvent fully evaporated. 10 μL of 98% sulfuric acid was then added to each sample and standard and the plate was once again incubated at 90°C for 2 minutes. The plate was placed on ice for five minutes after the incubation period and 190 μL of vanillin solution was added to each well. The plate was then shaken slowly in the plate reader for 15 minutes and then read at 525 nm. Absorbance values were recorded and used to calculate triglyceride concentration in mosquito samples.

### Water content

We used the protocol described in Benoit & Denlinger (2007) to determine water content in mosquitoes. Briefly, mosquitoes were collected on the day of their emergence, weighed at their initial mass, and then placed in a -20°C freezer for 6 hours to be killed. The samples were then transferred to an incubator (FisherBiotech Hybridization Incubator) set to 70°C. Daily measurements of each individual’s weight were made until the values became constant, indicating that there was no more water left in the samples. This final mass was then used to calculate the percentage of water within each individual.

### Estimates of *Wolbachia* density by qPCR

In both Cr and Fo mosquitoes, we determined the presence and density of *w*AlbA and *w*AlbB after temperature treatments (Dutton & Sinkins, 2004). Briefly, genomic DNA (gDNA) was extracted using the Wizard® Genomic DNA Purification Kit (Promega) from 16 larvae and 16 ovaries the day of emergence, 7 and 12-14 days post-emergence (dpe) for each thermal regime. Abundance of *w*AlbA and *w*AlbB was assessed in each sample by qPCR using the *Ae. albopictus* homothorax gene (AALC636_001297) as reference (Mancini et al., 2021). PCR reactions were performed in a total volume of 20 μl, containing 10 μl SYBR Green (ThermoFisher), 4 μl of gDNA, 4 μl of H_2_O and 1 μM of each primer: Forward (F) (GGGTTGATGTTGAAGGAG) and Reverse (R) (CACCAGCTTTTACTTGACC); F (AAGGAACCGAAGTTCATG) and R (AGTTGTGAGTAAAGTCCC); and F (TGGTCCTATATTGGCGAGCTA) and R (TCGTTTTTGCAAGAAGGTCA), for *w*AlbA-wsp, *w*AlbB-wsp, and the *Ae. albopictus* homothorax gene (AALC636_001297), respectively (Mancini et al., 2021; Martinez et al., 2022). The temperature was cycled at 95°C for 2 min, then 40 cycles at 95 °C for 5 seconds and 60 °C for 30 seconds, followed by the melting curve generation.

### Transcriptome analyses

We generated three larval pools, each consisting of ten 4th instar larvae randomly collected from each of the eight trays/thermal regime. We also collected females at the day of their emergence for a total of three pools of 10 mosquitoes each for each thermal regime. Larvae and adult females were sampled at the same time of the day. Both adult and larval samples were homogenised in 50 μL of Trizol (Life Technologies, Madrid, Spain) and stored at -80°C until RNA extraction. Total RNA was extracted using the standard Trizol protocol and re-suspended in 20 μl of nuclease-free water. Total RNA was sent to Macrogen Europe BV for quality control, TruSeq Stranded mRNA library preparation and sequencing on Illumina platform. Each library was sequenced pair-end (2 x 100 bp) at a depth of 30 million reads.

We used the nf-core/RNAseq bioinformatics pipeline (https://nf-co.re/rnaseq), which includes quality check and trimming of the reads for RNAseq analysis, using the AalbF2 genome assembly (Palatini et al., 2020). Fragments per kilobase of exon per Million mapped reads (FPKM) was used as a proxy for gene expression (Zhao et al., 2020). Analysis of gene differential expression (DE) among conditions was conducted with DeSeq2 in RStudio (2022.02.3) using the function “DESeqDataSetFromMatrix” (Love et al., 2014). We selected as differentially expressed genes (DEGs), those genes resulting with a Log2FC value ≥ |2| and a p-value < 0.01 from each comparison. Comparisons included intra-population comparisons of samples for larvae or emerging adults reared at the different thermal regimes (e.g., Fo larvae 18°C *vs.* Fo larvae 28°C). We also assessed inter-population DE genes between Fo and Cr across the same developmental stages and thermal regime (e.g., Fo larvae 18°C *vs.* Cr larvae 18°C). For comparisons among populations, genes that were DE between the two strains at 28°C were not considered to derive the list of DE genes between populations at both 18°C and 32°C to avoid accounting for population-specific differences. Relative fold changes in gene expression between samples were determined as a ratio of each FPKM; among strain comparisons were always Fo *vs.* Cr unless otherwise stated. Gene Ontology (GO) functional assignment and gene enrichment of DE genes followed the strategy outlined by Lozada-Chavez et al., (2023). Briefly, we created a custom annotation database (i.e. org.albopictus.eg.db R package) from merged results with Blast2GO (Götz et al. 2008) of three functional approaches: (1) Gene Ontology (GO) annotations covering ∼63% of the AALFPA-proteome, as retrieved from VectorBase v55 (Amos et al., 2022); (2) a homologs BLAST search of the AALFPA-proteome against the NCBI Diptera nr database v5; (3) a functional homologs search with InterProScan v5 (Jones et al., 2014) against four protein-domain databases: Pfam v33.1 (Mistry et al. 2021), ProSiteProfiles v20.2 (Sigrist et al., 2013), SUPERFAMILY v2.0 (Wilson et al. 2009), and TIGRFAM v15.0 (Haft et al., 2003). After this, 80% of the AALFPA-proteome was annotated. A GO enrichment analysis for major GO categories was performed with our in-house org.Aalbopictus.eg.db and clusterProfiler v4.2.2 (Wu et al., 2021) to identify functional groups that were enriched in our sets of DE genes. P-values (p≤0.05) obtained with clusterProfiler were corrected for multiple tests with the Benjamini-Hochberg procedure, and the redundancy of enriched GO terms for each major GO classification was removed with simplify, both from clusterProfiler.

### Data Analyses

We derived fecundity from our wing length (WL) data based on the function ln(egg number)=0.79+1.4*WL, which accounts for the hyperallometric relationship between wing length and fecundity in mosquitoes (Armbruster & Hutchinson, 2002; Nørgaard et al., 2022). The Shapiro-Wilk test was used to assess normality in the distribution of data (Ghasemi & Zahediasl, 2012); parametric ANOVA or the nonparametric Kolmogorov-Smirnov, Mann Whitney or Kruskal-Wallis tests were selected accordingly (Ramachandran & Tsokos, 2020). Statistical differences were considered significant if p-value≤0.05. Differences between populations and across temperatures in egg hatching rate, egg hatching time, larval viability, LDT (larval developmental time) and PDT (pupal developmental time), egg-to-adult developmental time (hereafter called developmental speed), wing length, adult sex ratio and body mass were tested through two-way ANOVA, with Tukey’s multiple comparison test (Lee & Lee, 2018). We tested Pearson’s correlation between wing length and body mass data of males and females in R (R version 4.2.2; R Core Team, 2022) using the ggpubr package (Kassambara, 2018). Pupal viability, *Tp*, KDT, energy reserves, water content and *Wolbachia* density data were analysed between populations and sexes with the Mann Whitney test and within strain with Kruskal-Wallis test, with Dunn’s multiple comparison test. *Tp* assay data were further analysed by the distribution with the Kolmogorov–Smirnov test (Massey, 1951). Longevity data were used to extrapolate the median survival time and the hazard ratio, which is a measure of how rapidly each mosquito died, with the Mantel-Haenszel method (Muenz et al., 1977). Comparisons of mosquito longevity were carried out with the Kaplan-Meier survival analysis, Log-rank (Mantel-Cox) test (Kishore et al., 2010). From LDT and PDT, we calculated larval developmental rate (LDR) and pupal developmental rate (PDR) and derived their ratio (LDR/PDR). LDR, PDR and LDR/PDR datasets were studied with regression analyses on developmental temperature as previously shown (Folguera et al., 2010; Jarošík et al., 2004). We used Prism 8 (GraphPad) for all statistical analyses.

To further quantify the effect of developmental temperature, population or sex, and their combinations on studied fitness traits (except for longevity), we used two-way ANOVA and obtained F-ratio (F) and the relative p-value, together with the percentage of variation explained by each source. We further used Principal Component Analysis (PCA) on life history traits and energy reserves, to explore the traits most influenced by temperature and population differences, using the function “fviz_pca_biplot” from the factoextra package (R version 4.2.2) (Kassambara & Mundt, 2020).

To compare the overall thermal performance of each population, we draw a Thermal Performance Curve (TPC) including data of egg hatching rate, egg hatching time, LDR, PDR, developmental speed, larval and pupal viability, sex ratio, longevity, wing length and body mass following a previously reported standardisation procedure of MacLean et al., (2019), in which TPCs were drawn based developmental viability, development speed, and adult fecundity. Briefly, a value of 1 is assigned to the highest value for each trait, and the rest of the values are processed as ratio. Curves were designed using the cubic spline method, setting as extreme values 10.4 °C, the lowest developmental temperature registered for *Ae. albopictus* (Delatte et al., 2009) and the mean *KDT* resulting from our study (46°C).

## RESULTS

We hatched eggs of the *Ae. albopictus* native tropical Fo and invasive temperate Cr populations at 18°C and 32°C and followed their development, including adult longevity, in comparison to mosquitoes reared at 28°C. In the same populations/conditions for which we analysed fitness traits, we further compared heat resistance and *Tp,* measured water content and energy reserve in emerging adults and performed transcriptome analyses in both larvae and adults at the date of emergence. Finally, we assessed prevalence of *w*AlbA and *w*AlbB in 4th instar-larvae and ovaries of females of the two populations sampled at 0, 7 and 12-14 dpe through qPCR (Figure 1).

**Figure 1.**
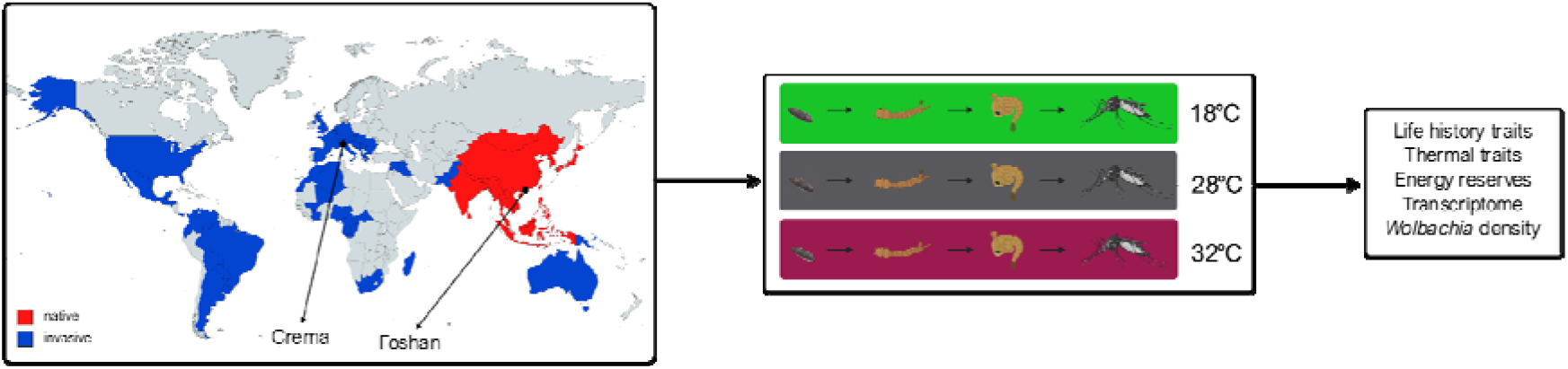
Experimental design: *Aedes albopictus* mosquitoes of the invasive Crema (Cr) and the native Foshan (Fo) populations were reared in the laboratory at three different temperatures (18, 28 and 32°C). For each condition, we measured life history traits (i.e. egg hatching rate, egg hatchability time, larval and pupal viability, larval and pupal developmental time, developmental speed, sex ratio, wing length, body mass, and adult longevity), thermal traits (thermal preference, and knock down temperature), energy reserves (i.e. water, protein, glycogen, triglycerides, and lipid contents), and *Wolbachia* density. Pools of larvae and adults were also sampled at emergence for transcriptome analyses. Created with BioRender and MapChart.

### Developmental temperature has a significant impact on mosquito fitness in a population-specific manner

While no temperature or population differences were observed in egg hatching rate, egg hatchability time was the lowest at 28°C in both Fo and Cr (Figure 2, Table S1). We also observed that when reared at 18°C, Cr mosquitoes had a significantly quicker hatching time than Fo ones (9.77±1.59 days in Cr, 14.01±3.70 days in Fo). A similar decreasing trend associated with higher developmental temperatures was observed in larval viability in Cr mosquitoes (larval viability was 70.99%±8.80 in mosquitoes reared at 18°C; 66.63%±7.47 at 28°C and 47.75%±8.12 at 32°C), while in Fo larval viability was not significantly altered by rearing mosquitoes under different thermal regimes (Figure 2, Table S1). In mosquitoes reared at 28°C, pupal viability was the lowest in Cr (pupal viability was 91.66±3.38, 87.93%±9.63 or 96.51%±3.94 in Cr mosquitoes reared at 18, 28 or 32°C, respectively), while the highest in Fo (pupal viability was 82.41±15.33, 89.52%±4.54 or 83.97%±10.10 in Fo mosquitoes reared at 18, 28 or 32°C, respectively). Developmental temperature also had significant effects on LDT, PDT, developmental speed, longevity and fecundity (Figure 2, Figure S2A, Table S1 and Table S2). Hatching eggs at 18 *vs.* 28°C increased the time for larval emergence by 4.6 times in Fo and almost 3 times in Cr. In both Cr and Fo, LDT and PDT were 2.5-3 times longer in mosquitoes reared at 18 *vs.* those reared at both 28°C and 32°C (Figure 2, Figure S2A, Table S1). Rearing mosquitoes at 18 *vs.* 28°C resulted in a decrease of almost 40% in developmental speed with no population differences (Table S1). Cr mosquitoes were larger than Fo mosquitoes at all tested temperatures, but wing length and body mass followed a similar trend in both populations with respect to developmental temperature (Table S1). Mosquitoes reared at 18°C were significantly larger than those reared at both 28°C and 32°C, except for Fo females that had their highest body mass (1.81±0.34 mg) when reared at 28°C (Figure 2). Through Pearson’s correlation, we showed that both wing size and body mass are significantly negatively correlated with temperature in both sexes (Figure 3, Table S3). Thus, wing length and body mass follow Bergmann’s rule, which states that in colder environments organisms are larger (Angilletta & Dunham, 2003).

**Figure 2.**
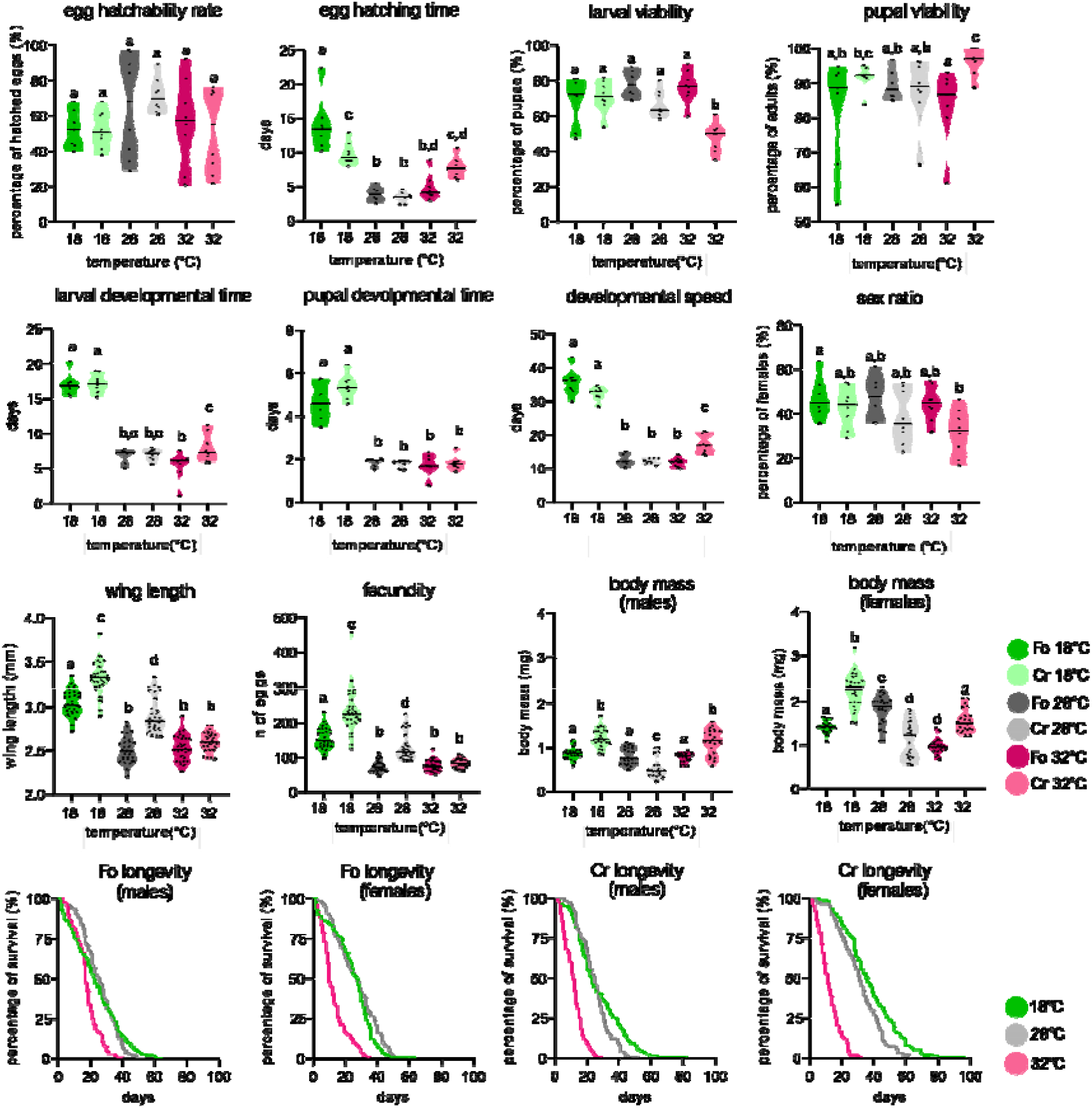
Life history traits of mosquitoes reared at 18°C (green), 28°C (gray) and 32°C (pink). Data for Fo and Cr mosquitoes are in darker or lighter colors, respectively. For each trait, solid lines in the violin plot represent the median value and data of each replicate or individual are shown as dots. Letters (“a” to “d”) refer to statistical comparisons: same letter denotes no significant difference (α >5%) between groups. Longevity is shown by survival curves for male and females, separately.

**Figure 3.**
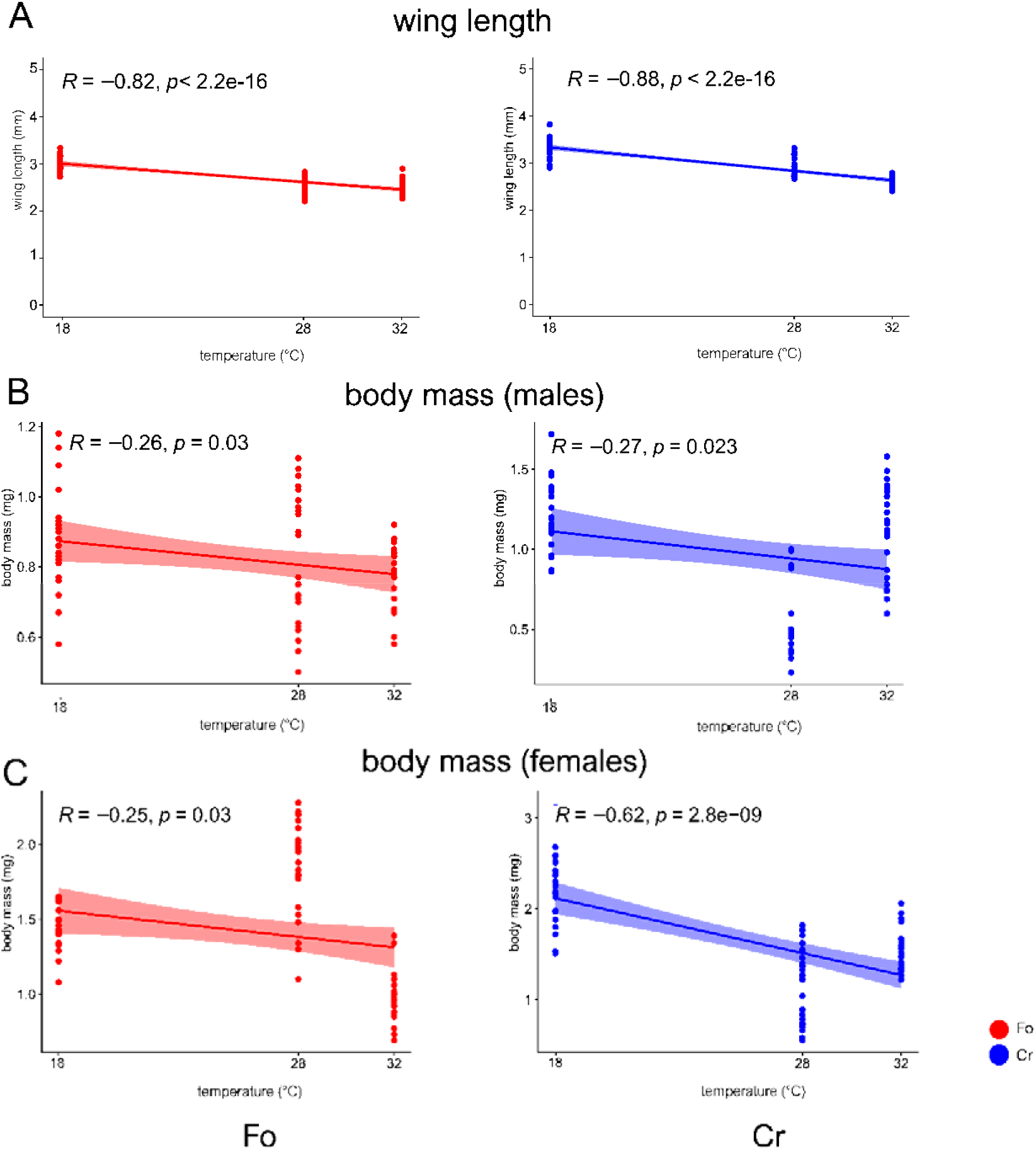
Pearson’s correlation between temperature and wing length **(A)** or body mass of males **(B)** and females **(C)**. Pearson’s correlation coefficient (R) is reported, together with the relative p-value. Fo and Cr data are shown in red and blue respectively.

Interestingly, the developmental temperatures across which changes in size were most observed differed between Fo and Cr mosquitoes: we saw a decrease of roughly 50% in body mass, and 15% in wing size, when Cr mosquitoes were reared at 18 *vs.* 28°C (body mass at 18°C: 1.21±0.21 mg in males, 2.23±0.38 in females; at 28°C: 0.53±0.22 mg in males, 1.12±0.42 in females; wing length at 18°C: 3.23±0.18 mm, at 28°C: 2.89±0.18 mm). This level of change in body mass was identified in Fo females when reared at 32°C with respect 28°C (body mass at 28°C: 1.81±0.34, at 32°C: 0.98±0.16 mg), while no temperature-related changes were observed in the body mass of Fo males (Table S1).

Developmental temperature also had an impact on adult longevity. In Cr, adult longevity was significantly reduced compared to mosquitoes reared at 18 *vs.* 28°C and to those reared at 28 *vs*. 32°C. However, in Fo significant differences were observed when comparing adult longevity of mosquitoes reared at both 18 or 28 *vs*. 32°C, but not between 18°C and 28°C (Figure 2, Table S1). In both Fo and Cr, the highest fecundity was observed in mosquitoes reared at 18°C (in Cr 233±61 eggs in Fo 154±30 eggs), compared to both 28°C (in Cr 124±36 eggs and in Fo 75±16 eggs) and 32°C (in Cr 84±12 eggs and in Fo 76±164 eggs), with Cr always exhibiting a higher fecundity compared to Fo mosquitoes (Table S1).

To verify these results were not stochastic, we re-performed fitness measurements in both Fo and Cr after rearing them in parallel at standard conditions for two years. We observed no statistical differences in any of the measured traits in neither Cr nor Fo (Figure S3).

Independently of the population and developmental temperature, mosquitoes had an average preferred temperature of 25.77 °C ± 6.62 and were knocked down at 46.5°C ±1.04, (Figure 4A and 4B). However, *Tp* patterns differed between Fo and Cr mosquitoes, with Cr females choosing a wider range of *Tp* values on the thermal gradient with respect to Fo females when reared at both 28°C and 32°C (*p* =0.0272) (Figure 4B).

**Figure 4.**
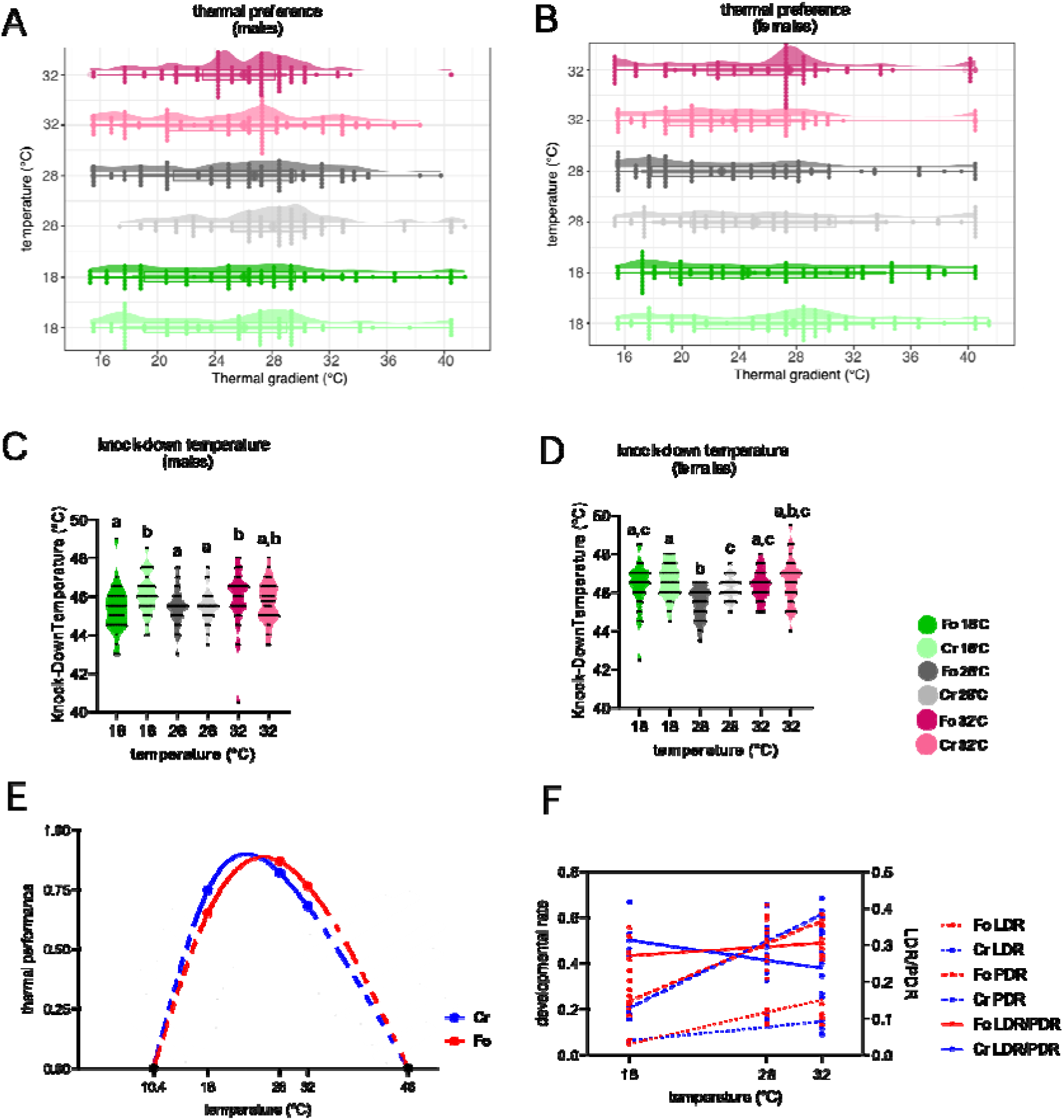
Raincloud plots of the *Tp* distributions for male **(A)** and females **(B)** and violin plots of the KDT of males **(C)** and **(D)** females reared at 18°C (green), 28°C (gray) and 32°C (pink). Data for Fo and Cr mosquitoes are in darker or lighter colours, respectively. Letters (“a” to “c”) refer to statistical comparisons: same letter denotes no significant difference between groups. Thermal performance curve of Fo (red) and Cr (blue) mosquitoes. Continuous lines represent the curves obtained through measured life history traits, whereas dotted lines are derived from estimated trait values **(E)**. Ontogenic trajectories of larval and pupal developmental time with respect to developmental temperature. The Y axis shows values of larval and pupal developmental rates on the left, their ratio (LDR/PDR) on the right **(F)**.

Using the measured KDT of 46.5°C (this study) and the critical lower thermal limit of 10.4°C for *Ae. albopictus* (Delatte et al., 2009), we built TPCs for both strains including data of egg hatching rate, egg hatching time, LDT and PDT, developmental speed, larval and pupal viability, sex ratio, longevity, wing length and body mass. We observed that both populations had a similar thermal breath, but a different thermal optimum which was 23.28°C for Cr mosquitoes and 25.78°C for Fo (Figure 4E).

Based on an ANOVA analysis, developmental temperature alone accounts for most of the variation in measurements of egg hatching rate, egg hatching time, LDT and PDT, development speed and wing length, whereas variation in measurements of larval viability is explained by the combination of developmental temperature and population (“temperature × population”) (Table S4). In summary, these results highlight that developmental temperature greatly affects *Ae. albopictus* life history traits with population differences.

### Testing developmental isomorphy at the population level

Because differences in developmental speed were significant between mosquitoes reared under different thermal regimes, we tested the hypothesis of developmental isomorphy, which is considered a general rule for ectotherms and states that developmental temperature does not alter the proportion of developmental time spent in a specific developmental stage (Jarošík et al., 2004). Regression analysis revealed a positive relationship between LDR and developmental temperature in both populations (Figure 4F). A similar trend was observed for PDR. We further compared the slope and the elevations of the LDR and PDR regressions on temperature within and between populations. In Cr, but not in Fo, LDR and PDR slopes are significantly different (Figure 4F, Table S5). No significant differences were found between populations. We further tested the regression on temperature of the ratio LDR/PDR, which showed isomorphy (b=0) in Fo, but a significantly negative slope (b=-0.005419) in Cr. These results indicate that in Cr temperate mosquitoes, temperature has specific effects on the duration of each life stage. On the contrary, Fo tropical mosquitoes do not change the proportion of time spent in the larval or pupal stage depending on the change in developmental temperature.

### Mosquito development at 18 and 32°C alters their energy reserves

Males showed comparable values of water content between populations, whereas Cr females had a higher water content than Fo ones when reared at both 28°C and 32°C (Figure 5). In both populations, water content was approximately 70% less in mosquitoes reared at 32°C with respect to those reared at both 18°C and 28°C (Figure 5, Table S6). A low content of water and water loss have been linked with reduced nutritional reserves in mosquitoes (Benoit et al., 2010), which aligns with what we observed here. We noted a lower content of proteins in both Fo and Cr mosquitoes reared at 32°C *vs.* those reared at 28°C and, in Cr mosquitoes, also a lower content of triglycerides and glycogen (Figure 5, Table S7).

**Figure 5.**
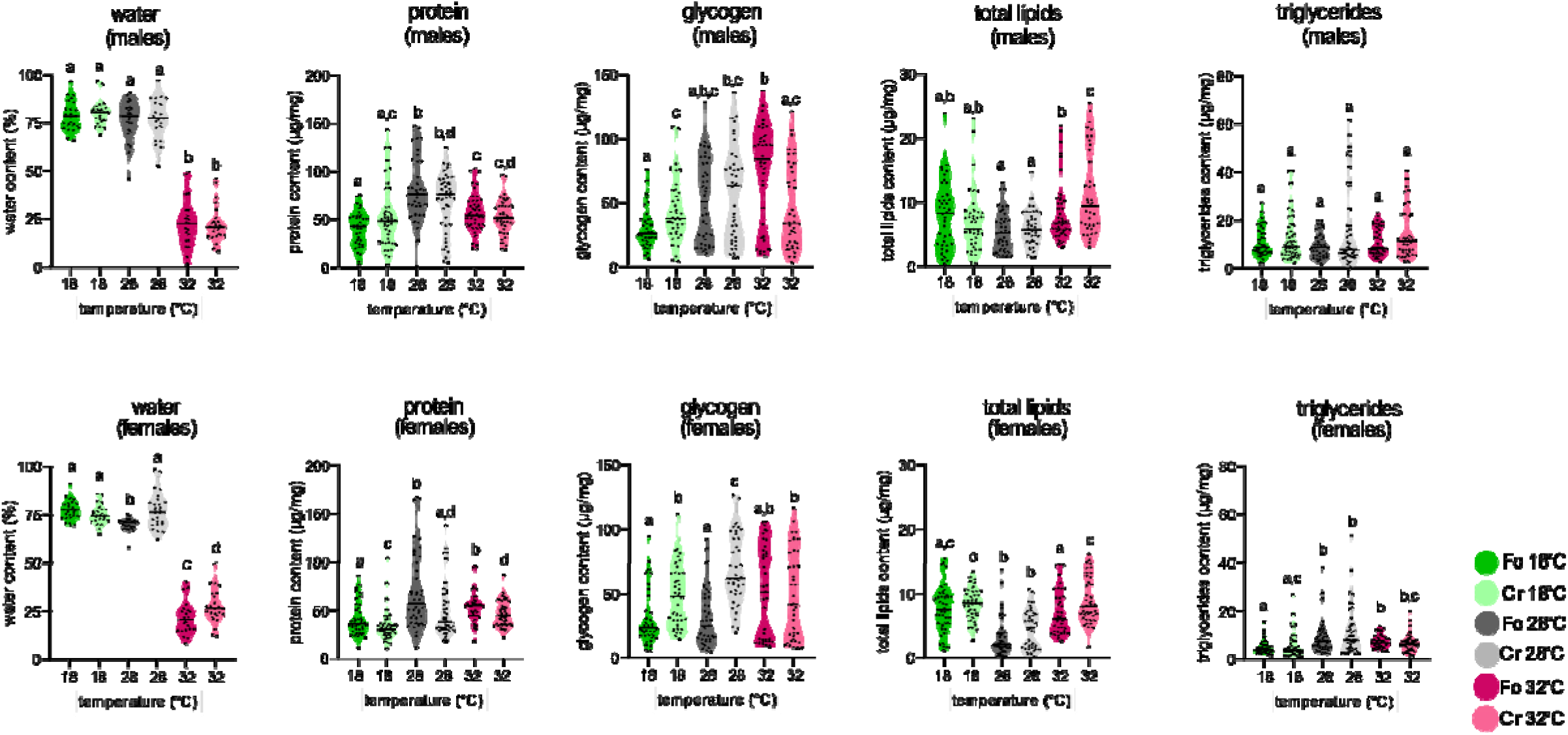
Water content and energy reserves of mosquitoes reared at 18°C (green), 28°C (gray) and 32°C (pink). Data for Fo and Cr mosquitoes are in darker or lighter colours, respectively. For each trait, solid lines in the violin plot represent the median value and data of each replicate or individual are shown as dots. Upper and lower plots are for males and females, respectively. Letters (“a” to “d”) refer to statistical comparisons: same letter denotes no significant difference between groups.

Overall, both reducing and increasing developmental temperature with respect to standard rearing condition had the same decreasing effect on protein and triglyceride content, with the exception of males where developmental temperature did not affect triglyceride content (Table S7). Sex differences in the accumulation of triglycerides are known in *Ae. aegypti* and could be linked to reproduction and egg maturation as triglycerides are involved in these processes (Arrese & Soulages, 2010; O’Meara & Van Handel, 1971). Glycogen content showed a similar trend as protein content in Cr; on the contrary, in Fo mosquitoes glycogen content increased with increasing developmental temperature, reaching its maximum value at 32°C (72.99 and 51.22 μg/mg, in males and females respectively). The opposite trend was observed in both populations for lipids, which significantly increased from their values in mosquitoes reared at 28°C (3.197±3.066 μg/mg in Fo; 4.373±3.050 in Cr μg/mg), reaching similar levels at l8 and 32°C (Table S7). PCA analyses confirmed that water and protein content were the nutrients mostly affected by temperature in both sexes (Figure S2B and S2C).

### More transcriptional changes in mosquitoes reared at 18°C than 32°C

Transcriptional analyses showed a higher number of DE genes in mosquitoes reared at 18°C than 32°C (for larvae: 155 DE genes at 18°C in Cr and 143 in Fo; at 32°C, 39 DE genes in Cr and 65 in Fo; for adults: 57 DE genes at 18°C in Cr and 236 in Fo; at 32°C, 26 DE genes in Cr and 33 in Fo) (Figure 6). Even though the identity of genes significantly DE at the different developmental temperatures was different, we found considerable overlap between populations in gene functions that were enriched in mosquitoes reared at either 18°C or 32°C with respect to those maintained at 28°C (Figure S4, Table S8). We saw an enrichment for lipid transporter activity and organonitrogen compound catabolic process in the transcriptome of adults reared at 18°C; hydrolase activity, acting on glycosyl bonds and ATP hydrolysis activity in adults reared at 32°C; chitin binding in larvae reared at 18°C and hydrolase activity acting on glycosyl bonds in larvae reared at 32°C (Table S8, Figure S4).

**Figure 6.**
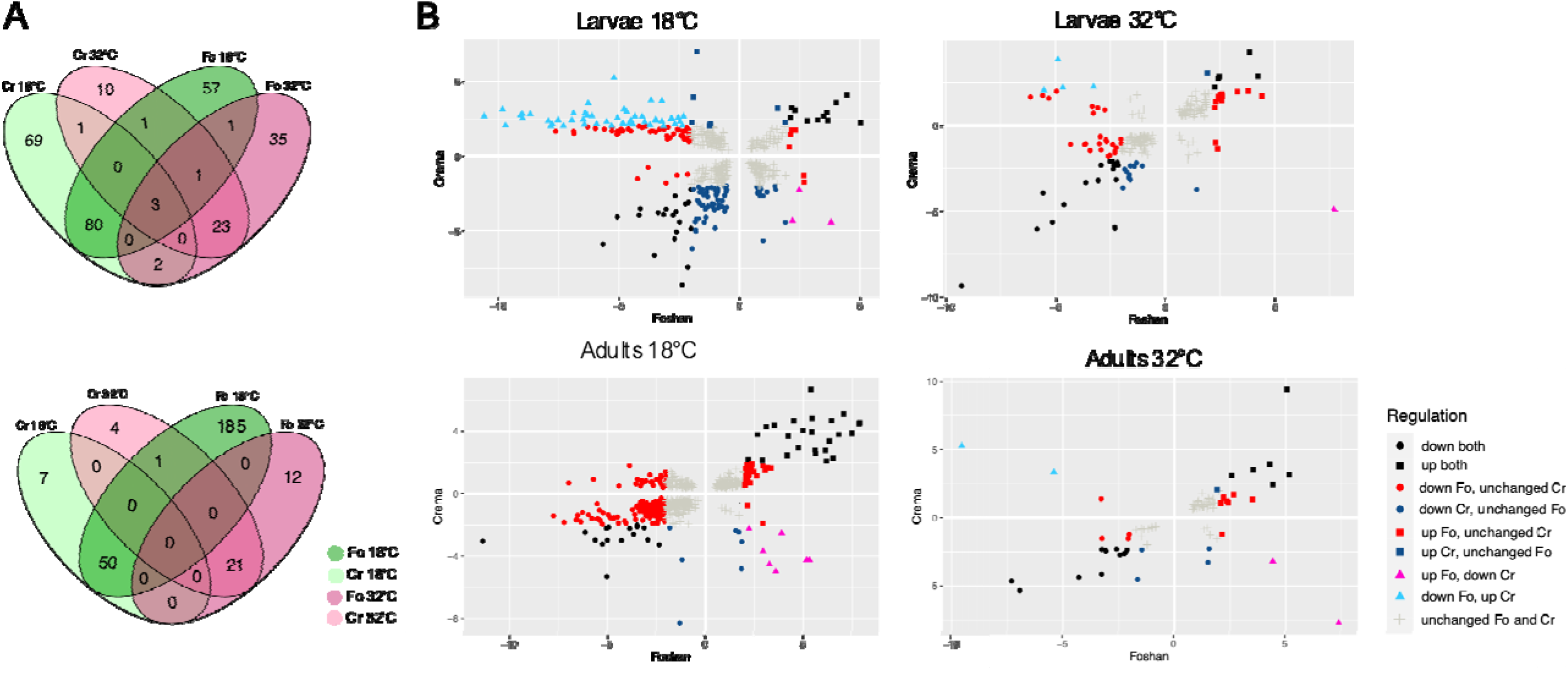
**(A)** Venn diagrams showing the number of DE genes across conditions as shown in the legend, in larvae (upper) and adult (lower). **(B)** Plots showing the differential expression of Fo and Cr mosquitoes at the larval (upper) and adult (lower) stages, when reared at either 18°C or 32°C. In each plot, a point indicates the expression level of a gene in Fo (x-axis) and Cr (y-axis) mosquitoes *vs*. their respective controls, meaning mosquitoes reared at 28°C. Differentially expressed genes are shown in different colours as shown by the legend; “unchanged” refer to genes that were not DE.

Given the high number of genes with unknown functions in the *Ae. albopictus* genome (Palatini et al., 2020), which could limit the resolution of a GO-based enrichment analysis, we further looked at candidate genes, including immunity-related genes, which are known to be modulated by temperature (Palatini et al., 2020; Wojda, 2017), and we compared our DE genes to candidate temperature-associated genes previously identified in *D. melanogaster* (Herrmann & Yampolsky, 2021). In adults of both populations, we saw a lower expression of cytochrome p450 genes (i.e. AALFPA_049004, AALFPA_074444, AALFPA_068814, AALFPA_046252, AALFPA_052377, AALFPA_080403), mostly in mosquitoes reared at 18 *vs.* 28°C, and heat shock proteins (i.e. AALFPA_060102, AALFPA_060249, AALFPA_041794, AALFPA_049001, AALFPA_067347, AALFPA_068981, AALFPA_051922, AALFPA_041503 and AALFPA_052248), mostly in mosquitoes reared at 32 *vs.* 28°C, in agreement with temperature-regulated genes previously identified in *D. melanogaster* (Table S9). We also observed the pickpocket-like gene (AALFPA_059066, AALFPA_080544) and genes related to muscle and cytoskeletal fluidity and movement (i.e. AALFPA_040928, AALFPA_051652, AALFPA_061734, AALFPA_053548, AALFPA_053957) to be upregulated in mosquitoes reared both at both 18 or 32 *vs.* 28°C (Figure 6) (Freundt & Linke, 2019; Litman & Stein, 2023; Nohria et al., 1992). Expression of immunity genes was regulated by temperature mainly at the larval stage, both in Cr and Fo, with similar functions (i.e. antimicrobial peptides, CLIP domain-containing serine proteases and fibrinogen-related proteins) (Palatini et al., 2020), but different genes, being elicited in the two populations (Table S10). Additionally, toll-like receptors (AALFPA_050924, AALFPA_052084, AALFPA_053911, AALFPA_053911 and AALFPA_063239) were upregulated up to 97.86 folds more in both adult and larvae reared at either 18 or 32 *vs.* 28°C in Cr mosquitoes, whereas in the Fo mosquitoes, the highest differential expression was identified for the antimicrobial peptide defensin-A (AALFPA_065006), which was upregulated 8.87 times in adults reared at 32°C with respect to mosquitoes reared at 28°C, and spaetzle 4 (AALFPA_059265), which was downregulated 62.28 and 9.24 in larvae reared at 32°C or 18°C, respectively (Table S10).

### Development at 18°C restricts *Wolbachia* density in *Ae. albopictus*

At the larval stage, Cr mosquitoes showed a higher concentration of both *wAlbA* and *wAlbB* than Fo mosquitoes at all developmental temperatures (Table S11). Additionally, *wAlbB* density was highest in Cr larvae reared at 18°C than in those reared at either 28°C or 32°C (Figure 7). This trend was maintained in ovaries of emerging females and extended also for *wAlbA*. In ovaries of mosquitoes sampled 7 and 12-14 dpe, the density of both strains of *Wolbachia* increased in mosquitoes reared at 28°C and 32°C, but not 18°C. Overall, these results show that developmental temperature influences *Wolbachia* density, with 18°C being a limiting temperature for both strains of *Wolbachia*. At all tested thermal developmental regimes, we observed a higher density of w*AlbB* than w*AlbA* than previously reported (Hu et al., 2020)

**Figure 7.**
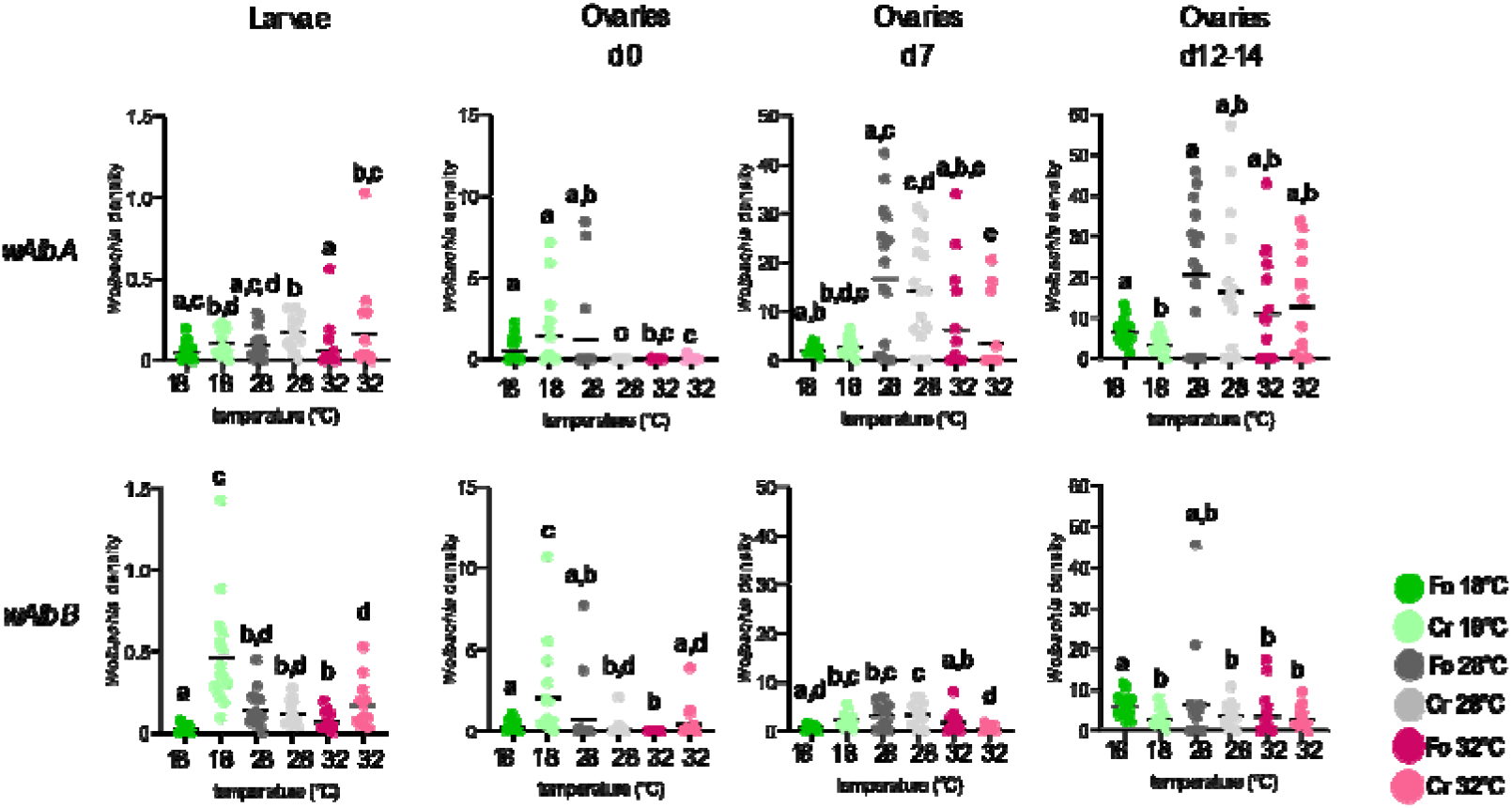
Density of *wAlbA* and *wAlbB* in Fo and Cr larvae and ovaries sampled at emergence (d0), 7 days post-emergence (d7) and 12-14 days after emergence (d12-14). Data for mosquitoes reared at 18°C, 28°C and 32°C are in green, grey and pink, respectively. Data for Fo and Cr mosquitoes are in darker or lighter colours, respectively. Letters (“a” to “e”) refer to statistical comparisons: same letter denotes no significant difference between groups.

## DISCUSSION

Anthropogenic climate change is rapidly changing natural environments favouring the expansion of invasive and ecologically plastic insect species, and advancing their phenology. Understanding how mosquito vector species respond to these changes is paramount not only to predict their seasonal activities and distribution range, but also to tailor effective control strategies. Here, we compared the effects of developmental temperature on mosquitoes from a temperate and a tropical population of the arboviral vector *Ae. albopictus*. We showed that, in this species, fecundity, egg hatching rate, egg hatching time, body mass and wing length, larval viability, LDT, PDT, developmental speed, and adult longevity are strongly influenced by developmental temperature with significant differences between populations. When reared at 18°C, Cr temperate mosquitoes had significantly higher fecundity and a longer longevity in females than tropical Fo ones, suggesting an overall better vector fitness performance. Furthermore, developing at 18°C was not a limiting factor for the accumulation of energy reserve in Cr mosquitoes as content of glycogen and protein in emerging adults was higher than that observed in Fo mosquitoes. This result was coupled with an upregulation of genes encoding for constituents of cuticle and carbohydrate metabolism in larvae, and lipid transporter and endopeptidase activities in adult mosquitoes. Chitin is one of the major components of insect cuticle, which is already known to be a dynamic structure, involved in protecting against water loss and mechanical damage (Muthukrishnan et al., 2020), and it is shaped by environmental conditions (Rajpurohit et al., 2021). Furthermore, chitin is also a constituent of the insect peritrophic membrane (PM), which is involved in food digestion and in protecting the epithelium against mechanical damage caused by food (abrasion) and against microorganisms (Terra, 2001). Endopeptidases are digestive enzymes, which are expressed primarily in the mosquito gut and are involved in the digestion of proteins of the blood meal (Borovsky, 2003). Glycogen can be mobilised in response to thermal stress (Storey, 1997) and is used to produce trehalose and sugar alcohols (polyols) that prevent cellular damage at low temperatures (Danks et al., 1994) by protecting membranes and maintain protein stability (Salvucci, 2000). Stored lipids and glycerol are processed and utilised as cryoprotectants (Chowanski et al., 2015).

In both populations, vector fitness performance, resulting from larval viability, adult longevity and fecundity, was lowest in mosquitoes reared at 32°C. In Cr, we also observed reduced larval viability, but increase in LDT, PDT, and egg hatching time with respect to mosquitoes reared at 28°C, suggesting that development at 32°C is more restrictive. This result correlates with a lower content of water, fat, and glycogen, a trend which has been regarded as a bioenergetic marker for stressful conditions (Sokolova, 2013). Additionally, in mosquitoes reared at 32°C, we also observed upregulation of genes associated with ion transport and binding, along with a 27.955 fold increase of the pickpocket protein 28-like gene (AALFPA_059066), which in *D. melanogaster* is essential for behavioural responses to noxious heat (Zhong et al., 2010) in Cr mosquitoes, and a downregulation of cuticle proteins (i.e. AALFPA_048721, AALFPA_071760 and AALFPA_079957 in Cr and AALFPA_041580 and AALFPA_063074 in Fo), which have been previously associated with desiccation stress in *D. melanogaster* (Horváth et al., 2023) and of heat shock proteins (*HSP*). In *Ae. aegypti,* higher thermal tolerance has been associated with lower expression of *HSP* genes suggesting expression of heat-shock proteins is expensive and may cause toxicity at the cellular level (Ware-Gilmore et al., 2023). Over-expression of genes associated with ion transport and binding correlates with heat causing alterations in pH and ion concentrations, that affect macromolecule functions and structure, and result in cellular damages (Denlinger & Yocum, 1998). These results are aligned with previous analyses of the early stages of development of *Ae. albopictus* mosquitoes from Brazil and La Réunion Island, which showed that 35°C is a limiting temperature for the development of larvae and the viability of pupae (Delatte et al., 2009; Monteiro et al., 2007). Thus, when environmental conditions result in an effective developmental temperature equal or above 32°C, the fitness performance of *Ae. albopictus* emerging in these conditions could be reduced. Our data on thermal preference show that, when reared at 32°C, mosquitoes choose temperatures between 24.21-26.10°C (Table S1). Consequently, the existence of suitable microenvironments with an *Ae. albopictus*-favourable T*_a_* and the levels of humidity will also have to be considered to precisely predict species distribution and abundance locally. Interestingly, temperatures between 24.09-26.93°C were also chosen by mosquitoes that had been reared at both 18°C and 28°C with no population difference.

The question arises about what these results mean in relation to the presence of *Ae. albopictus* in temperate regions of the world as the Mediterranean basin. The Mediterranean basin is considered a climate change hotspot, with an increase of about 1.54°C in the average T*_a_* since 1980, and also an increase in the minimum and maximum temperature extremes (Fioravanti et al., 2016; Toreti & Desiato, 2008), that translate into a net increase of about 14% of the number of summer days, an increase in the number of tropical nights (defined as nights with a minimum temperature exceeding 20°C) and hot days (defined as days with a maximum temperature above 35°C) and a warming trend in all seasons (Fioravanti et al., 2016). Under this scenario, our results suggest that *Ae. Albopictus* fitness performance would be at its highest when mosquito population emerges in late Spring (source: World-weather, https://world-weather.info/forecast/italy/rome/july-2023/). These data further highlight the higher risk for transmission of chikungunya virus (CHIKV) with respect to DENV in Europe given that CHIKV transmission is supported in *Ae. albopictus* at 20°C, a temperature that is not permissive for DENV (Mercier et al., 2022). Importantly, our experimental design measured changes in mosquitoes directly exposed to a new developmental temperature; we cannot exclude that long-term (over generations) exposure to 18°C or 32°C could result in thermal adaptation that manifests in different fitness and physiological responses as those observed here during thermal acclimation (Lagerspetz, 2006).

We also did not observe strain differences in TPCs, contrary to the expectation that insects from temperate climates, which experience a larger range in temperatures across the year than those from tropical climates, are more plastic and have a broader thermal sensitivity than tropical insects (Angilletta, 2009; Huey & Kingsolver, 1989). The absence of this trend in our data is consistent with previous results (Mac Lean et al., 2019; Tippelt et al., 2020), and may be related to the recent, rapid and chaotic global dispersal of *Ae. albopictus,* which overlaid populations of different geographic origins in temperate Europe (Manni et al., 2017). Nonetheless, our data clearly showed that temperate Cr mosquitoes exhibit their best fitness performance at a lower temperature than tropical ones. Importantly, measured fitness traits are maintained after rearing the two populations in parallel for two years in laboratory, suggesting thermal adaptation, and indicating that detected differences will possibly magnify under thermal regimes close to the lower thermal limit of *Ae. albopictus*, estimated at 10.4°C (Delatte et al., 2009). This conclusion is further supported by the observation that while tropical Fo mosquitoes follow developmental isomorphy, temperature had a stage specific effect on the duration of the larval and pupal stages in Cr mosquitoes. Lack of isomorphy in Cr mosquitoes was mainly driven by the PDR of mosquitoes reared at 18°C, with a longer PDT resulting in the emergence of adults with a high energy reserve content, not significantly different than that of mosquitoes developing at 28°C. Longer LDT and PDT in mosquitoes reared 18°C would favour interventions targeting juvenile stages, which have been shown to be more susceptible to thermal stress than adults possibly due to their dependence on ephemeral environments, with no possibility to move away (Reinhold et al., 2018).

Finally, our data showed that constant exposure to 18°C during mosquito developmental stages induced a decrease in the density of the native *Wolbachia* strains, *wAlbA* and *wAlbB*, compared to those reared at 28°C and 32°C. On the contrary, exposure at 32°C did not represent a thermal stress for native *Wolbachia.* Similarly, exposure to 18°C during development resulted in a drop in density of other native *Wolbachia* strains in *D. melanogaster* and in *Nasonia vitrippenis* (Bordenstein & Bordenstein, 2011; Chrostek et al., 2021). This further suggests that temperature likely shapes *Wolbachia*-host interactions in nature and, as a result, frequencies of symbionts in different host populations. However, studies using diurnal field relevant temperature cycles, and possibly other ecological parameters, should be considered using different host populations and *Wolbachia* variants. These findings reinforce the importance of considering abiotic factors in predicting symbiont stability, and in turn, in providing a comprehensive understanding of the repercussions of increased environmental temperatures on insect vectors.

## ACKNOWLEDGEMENTS

We are grateful to José E. Crespo and Claudio Lazzari for providing the device used in the present study to measure the *KDT*.

## AUTHOR CONTRIBUTION

M. Carlassara performed biological experiments, analyzed the data and wrote the manuscript; A. Khorramnejad contributed in fitness analyses and analyzed the data; H. Oker performed energy reserves and contributed in thermal trait experiments; R. Bahrami contributed to fitness experiments; A. N. Lozada-Chavez contributed with bioinformatic analyses of transcriptomic data and statistical analyses of the data; M.V. Mancini contributed in experiments of *Wolbachia* prevalence; M. Body contributed in energy reserve experiments and data analyses; C. Lahondère conceived the study, performed analyses of thermal traits and energy reserved, analyzed data and wrote the manuscript; M. Bonizzoni conceived and supervised the study, obtained funding, wrote the manuscript with feedback from all authors.

## SUPPLEMENTARY MATERIAL

**Table S1.** Comparison of values of life-table parameters and thermal traits determined for each population at different developmental temperatures (A-comparisons within population) and at each developmental temperature between population (B-comparisons between populations)

**Table S2.** Comparison of adult longevity for each strain and sex at different developmental temperatures (A-comparisons within population) and at each developmental temperature between populations (B-comparisons between populations)

**Table S3.** Pearson’s correlations between given traits (wing length and body weight) and temperature.

**Table S4.** Effects of developmental temperature, population, sex and their combinations on fitness traits (two-way and three-way ANOVA). Degrees of freedom (df), Mean Squares (MS), F ratio (F), p-value (p) and the percentage of variation (% of total variation) are reported.

**Table S5.** Developmental isomorphy. Regression analysis report with b (slope) and the relative p-value (A) and the relative p-value for each comparison in slope and elevation (B).

**Table S6.** Comparison of water content for each strain and sex at different developmental temperatures (A-comparisons within population), at each developmental temperature between populations (B-comparisons between populations) and at a set developmental temperature and population between sexes (C-comparisons between females and males)

**Table S7.** Comparison of energy reserves, meaning content of protein, glycogen, lipid and triglycerides, for each population and sex at different developmental temperatures (A-comparisons within population), at each developmental temperature between populations (B-comparisons between populations) and at a set developmental temperature and population between sexes (C-comparisons between females and males)

**Table S8.** GO enrichment of genes differentially expressed in each population at either 18°C or 32°C with respect to expression at 28°C; genes differentially expressed between Fo and Cr at 28 were excluded. Molecular Function, GO ID, adjust p-value (p.adjust), genes involved, Gene Ratio (ratio of the number of genes assigned to the specific GO and the total number of molecular function-annotated genes present in the given list of DE genes), Background Ratio(BgRatio, ratio of the number of genes assigned to that specific GO and the total number of genes present in the genome which are molecular-function annotated), Fold Enrichment (FE) and Regulation are reported for each strain, life stage and temperature.

**Table S9**. Genes identified as DE in our transcriptomic analyses and responsive to temperature also in *D. melanogaster* (Herrmann & Yampolsky, 2021)

**Table S10**. List of immunity genes that were differentially expressed in each population at either 18°C or 32°C with respect to expression at 28°C; genes differentially expressed between Fo and Cr at 28°C were excluded.

**Table S11.** *Wolbachia* relative abundance (qHTH/qwAlb). Kruskal-Wallis test within strain, Mann-Whitney test between populations. P-values are reported for each comparison.

**Figure S1.**
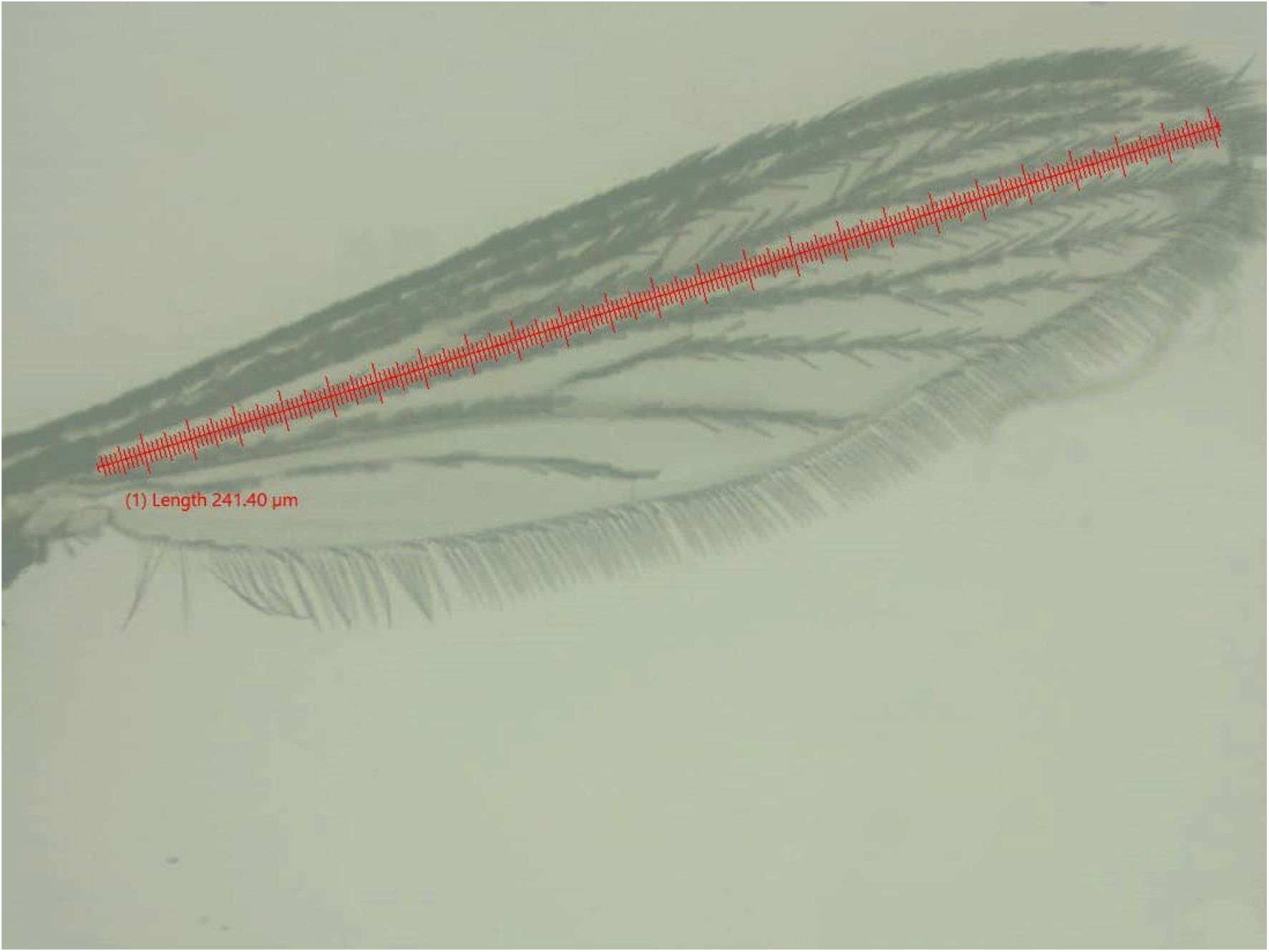
Dissected right wing was measured from the axial incision to the apical margin, excluding the fringe of the scales (magnification 20x) as shown by the red scale.

**Figure S2.**
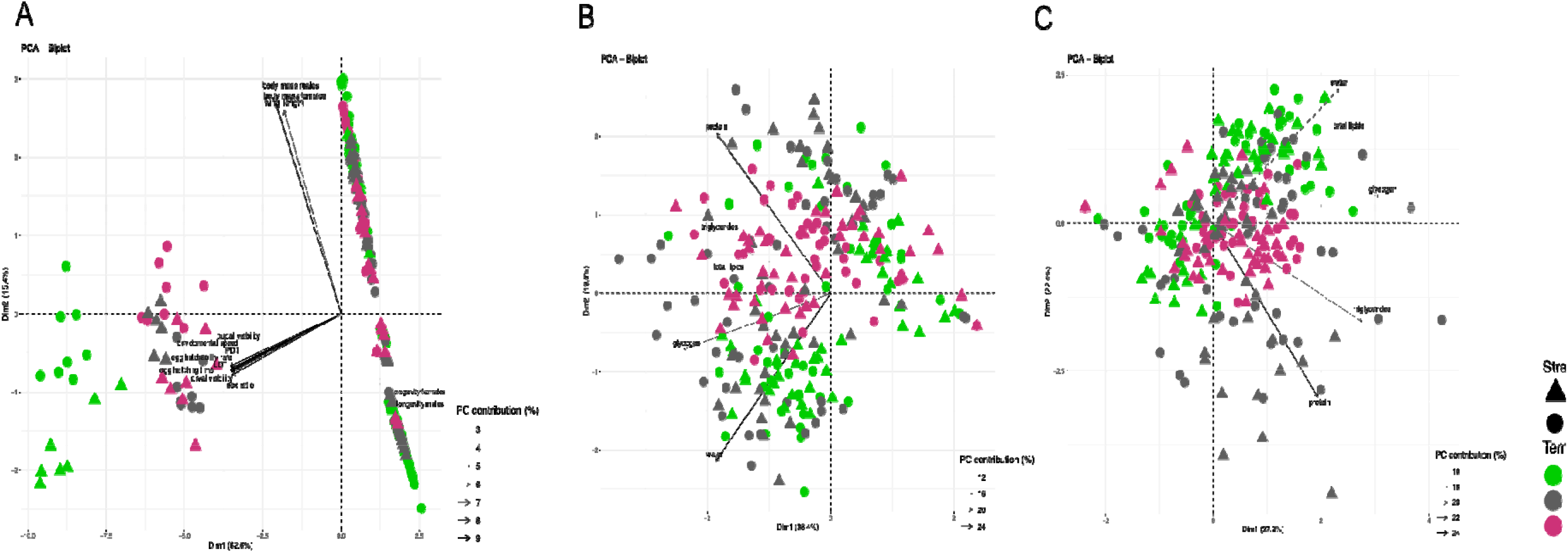
Principal Component Analysis showing the contribution of life history traits **(A)** and energy reserve in males **(B)** and females **(C)** of Fo (triangle) and Cr (circle). Data from mosquitoes reared at 18°C, 28°C, and 32°C are in green, gray, and pink, respectively.

**Figure S3.**
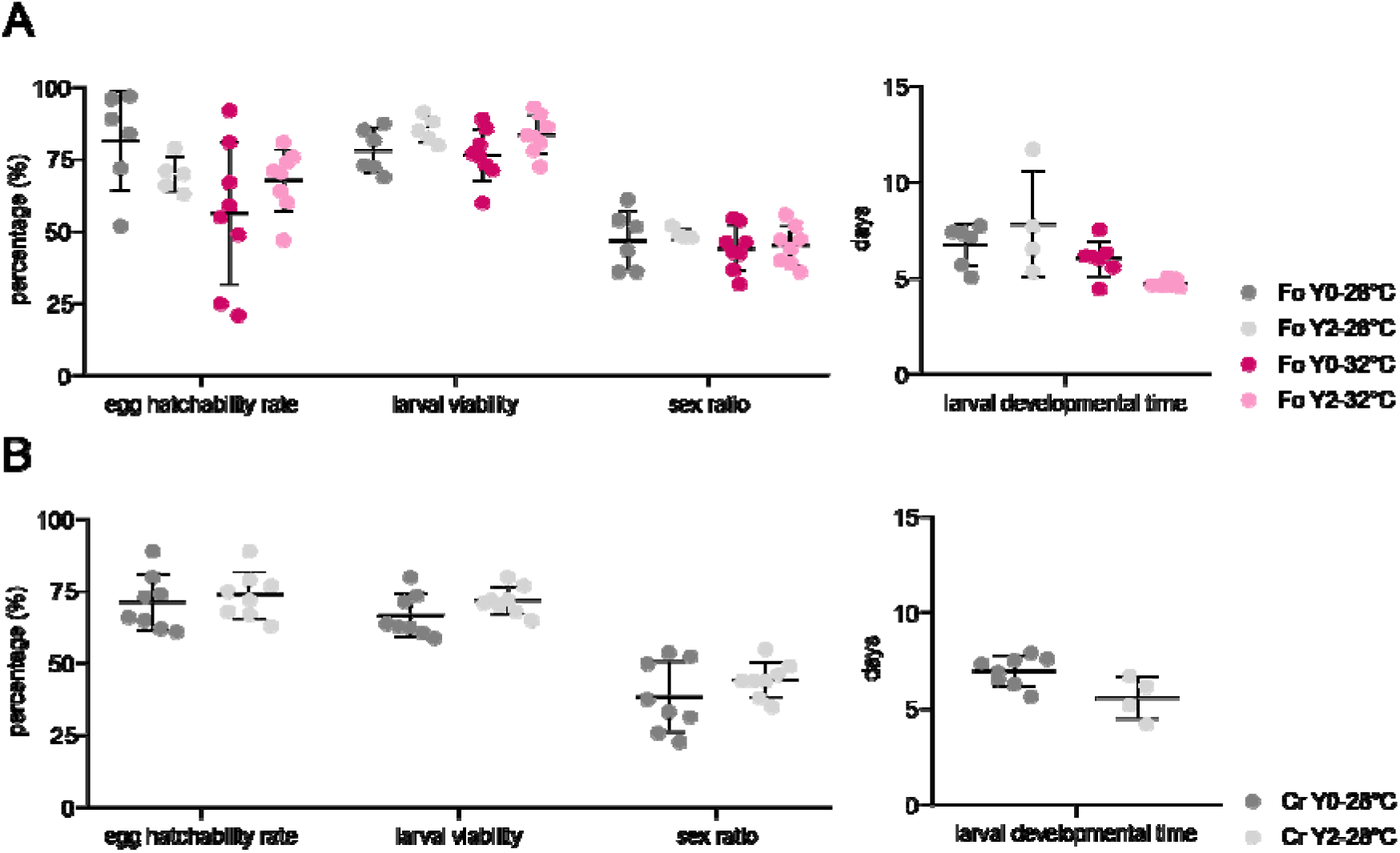
**(A)** Comparison of life history traits values as estimated in Fo mosquitoes after a two-year span (Y0 *vs.* Y2). Data were measured in mosquitoes reared at 28°C (grey dots) and 32°C (pink dots). **(B)** Comparison of fitness traits of Cr mosquitoes reared at 28°C after a two-year span (Y0 *vs*. Y2).

**Figure S4.**
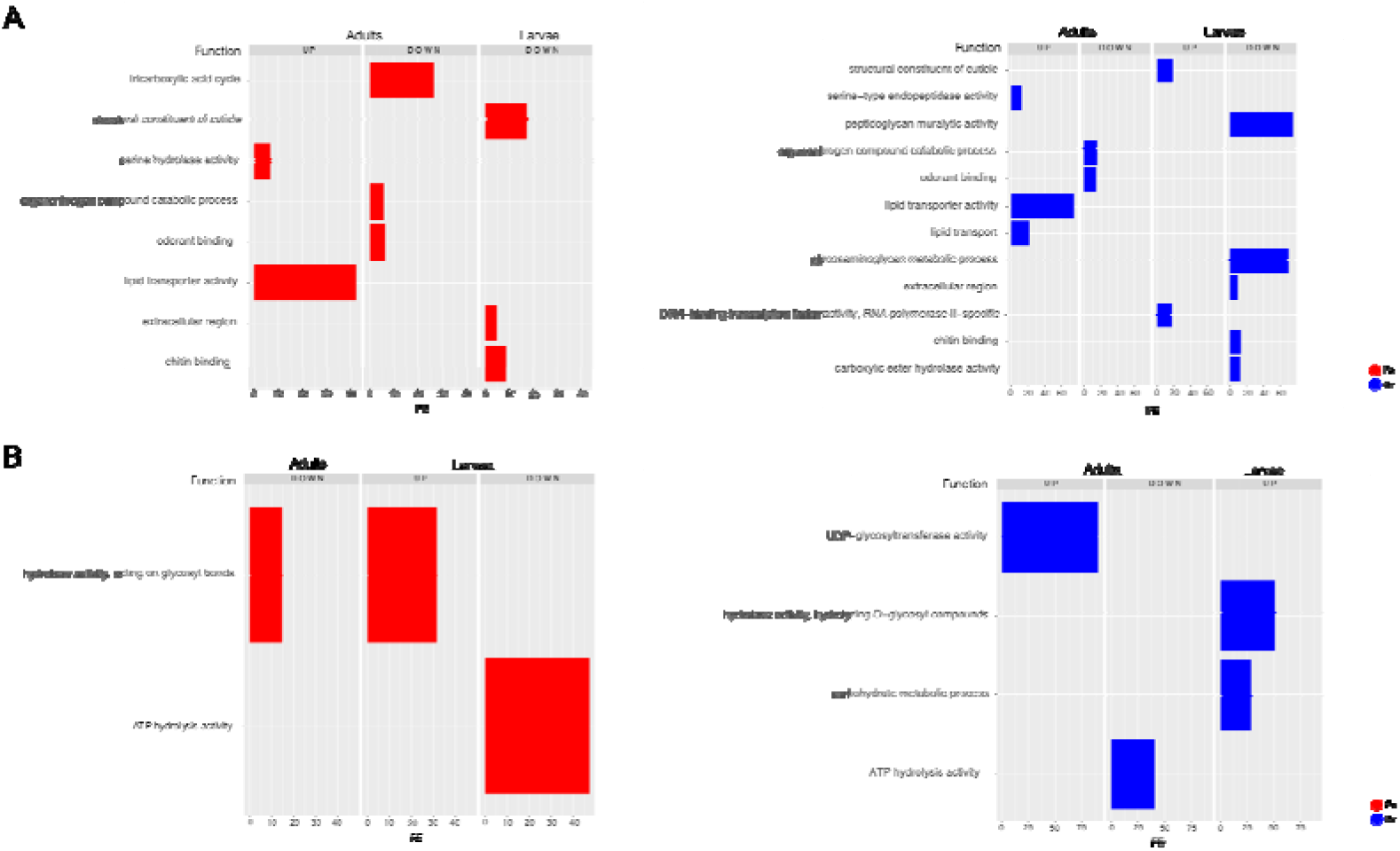
GO enrichment profiles reporting the fold-enrichment (FE) are compared between populations in adults and larvae of mosquitoes reared at 18°C **(A)** and 32°C **(B)**. Data for Fo and Cr are in red and blue, respectively.

